# Canine cognitive dysfunction (CCD) patients have reduced total hippocampal volume compared with aging control dogs: a comparative MRI study

**DOI:** 10.1101/2020.01.27.920918

**Authors:** Curtis Wells Dewey, Mark Rishniw, Simon Platt, Kelsey Robinson, Joseph Sackman, Marissa O’Donnell

**Author notes:** AUTHOR CONTACT INFORMATION: CW Dewey: C4 169 Clinical Programs Center, Cornell University College of Veterinary Medicine, Ithaca, NY 14853. M Rishniw: C2 015 Clinical Programs Center, Cornell University College of Veterinary Medicine, Ithaca, NY 14853. SR Platt: Department of Small Animal Medicine & Surgery, College of Veterinary Medicine, University of Georgia, 2200 College Station Rd, Athens, GA 30602. K Robinson: Department of Small Animal Medicine & Surgery, College of Veterinary Medicine, University of Georgia, 2200 College Station Rd, Athens, GA 30602. J Sackman: Long Island Veterinary Specialists, 163 South Service Road, Plainview, NY 11803;. M O’Donnell: Long Island Veterinary Specialists, 163 South Service Road, Plainview, NY 11803;.

## Abstract

Hippocampal atrophy is a key pathologic and MRI feature of human Alzheimer’s disease (AD). Hippocampal atrophy has not been documented via MRI in canine cognitive dysfunction (CCD), which is considered the dog model of human AD. The purpose of this retrospective comparative volumetric MRI study was to compare total hippocampal volumes between successfully aging (control) dogs and dogs diagnosed with CCD. Mimics^®^ software was used to derive total hippocampal volumes and total brain volumes from the MRI studies of 42 aging dogs (≥ 9 years): 16 dogs diagnosed with CCD and 26 successfully aging controls. Total hippocampal volume normalized to total brain volume was significantly less for CCD patients compared with control dogs (p=0.04). The results of this study suggest that-similar to human AD-hippocampal atrophy is a pathological feature of CCD. This finding has potential importance for both investigating disease mechanisms related to dementia as well as future hippocampal-targeted therapies.

## Introduction

Alzheimer’s disease, a degenerative brain disorder of people, shares many clinical and pathological features with canine cognitive dysfunction (CCD), a disorder affecting aging dogs. Consequently, investigators consider CCD a naturally occurring model for studying human Alzheimer’s disease. Furthermore, CCD commonly causes frustration for dog owners and veterinarians.^1,2^ Studies have documented hippocampal damage as an early and prominent pathologic feature in both Alzheimer’s disease and CCD and attributed this pathology to the deposition of neurotoxic compounds such as beta-amyloid and tau proteins.^3-5^

In humans, volumetric measurement of the hippocampi from magnetic resonance images (MRI) allows clinicians to assess the presence or absence of hippocampal atrophy, as a diagnostic marker for Alzheimer’s disease.^6-8^ Additionally, MRI-based hippocampal volumetric measurements are being evaluated as means of assessing responses to future hippocampal-based treatment options for Alzheimer’s disease.^8,9^

The use of MRI to assess hippocampal volume in dogs with CCD has not been reported. The purpose of this MR imaging study was to compare total hippocampal volumes between dogs with CCD and similarly aged control dogs. We hypothesized that dogs with CCD would have smaller total hippocampal volumes compared with controls.

## Materials and Methods

We searched MRI databases from five institutions (Cornell University Hospital for Animals, University of Georgia, Long Island Veterinary Specialists, Veterinary Specialty and Emergency Services of Rochester and Oradell Animal Hospital) for brain MRI scans of aging (≥ 9 yrs old) dogs diagnosed with CCD and similarly aged dogs with no evidence of CCD (controls). Control dogs had undergone MRI imaging for reasons unrelated to CCD, including peripheral vestibular dysfunction, late-onset epilepsy, Horner’s syndrome and blindness. Because relatively few aged dogs undergo cranial MRI scans for non-CNS disorders, we expanded our control group by acquiring additional control MRI scans from two sources: 10 mixed-breed retired sled dogs with normal neurologic examinations that had been imaged as part of another study and 6 neurologically normal small-breed dogs whose owners volunteered for a no-cost brain MRI prior to scheduled dentistry procedures. Because of the nature of this study, the need for IACUC approval was waived by Cornell University’s Institute for Animal Care and Use Committee.

We based our diagnosis of CCD on previously established historical and clinical criteria together with characteristic MRI abnormalities (excluding hippocampal measurements).^1,10,11^ In addition, we only included cases of CCD for which this diagnosis was clearly stated in the medical record and supported by the MRI report.

All MRIs were performed under general anesthesia with one of six magnets: 1) 1.5 T Siemens Avanto (Munich, Germany) 2) 1.5 T Toshiba Vantage Elan (Lake Forest CA, USA) 3) 3.0 T Philips Achieva (Nutley NJ, USA) or 4) 3.0 T GE Discovery MR750 (Chicago IL, USA). Imaging sequences acquired included the following: sagittal T2-weighted; transverse T2-and T1-weighted; transverse and dorsal plane T1-weighted post-gadolinium injection; transverse T2-fluid attenuated inversion recovery (FLAIR); and transverse T2* gradient-recalled echo (GRE). For the 1.5-T MRI units, measurement parameters were as follows: slice thickness, 3.5 mm; slice gap, 3.5 mm; FOV, 185 mm; matrix size of images, 480 × 480. For the 3.0-T MRI units, measurement parameters were as follows: slice thickness, 2.0 mm; slice gap, 1.0-3.0 mm (depending on dog size); FOV, 1101 mm; matrix size of images, 400 × 400.

For each dog, three-dimensional volumes were measured from T2-weighted brain images using Mimics^®^ software by two observers (JS and MO) who were unaware of the status of the dogs in the study. Quantitative volumetric measurements of both hippocampi as well as total brain volume were acquired for each dog, as previously described (Figure 1).^12^ Anatomic landmarks for measurements were used from published reference information.^13,14^

**Figure 1.**
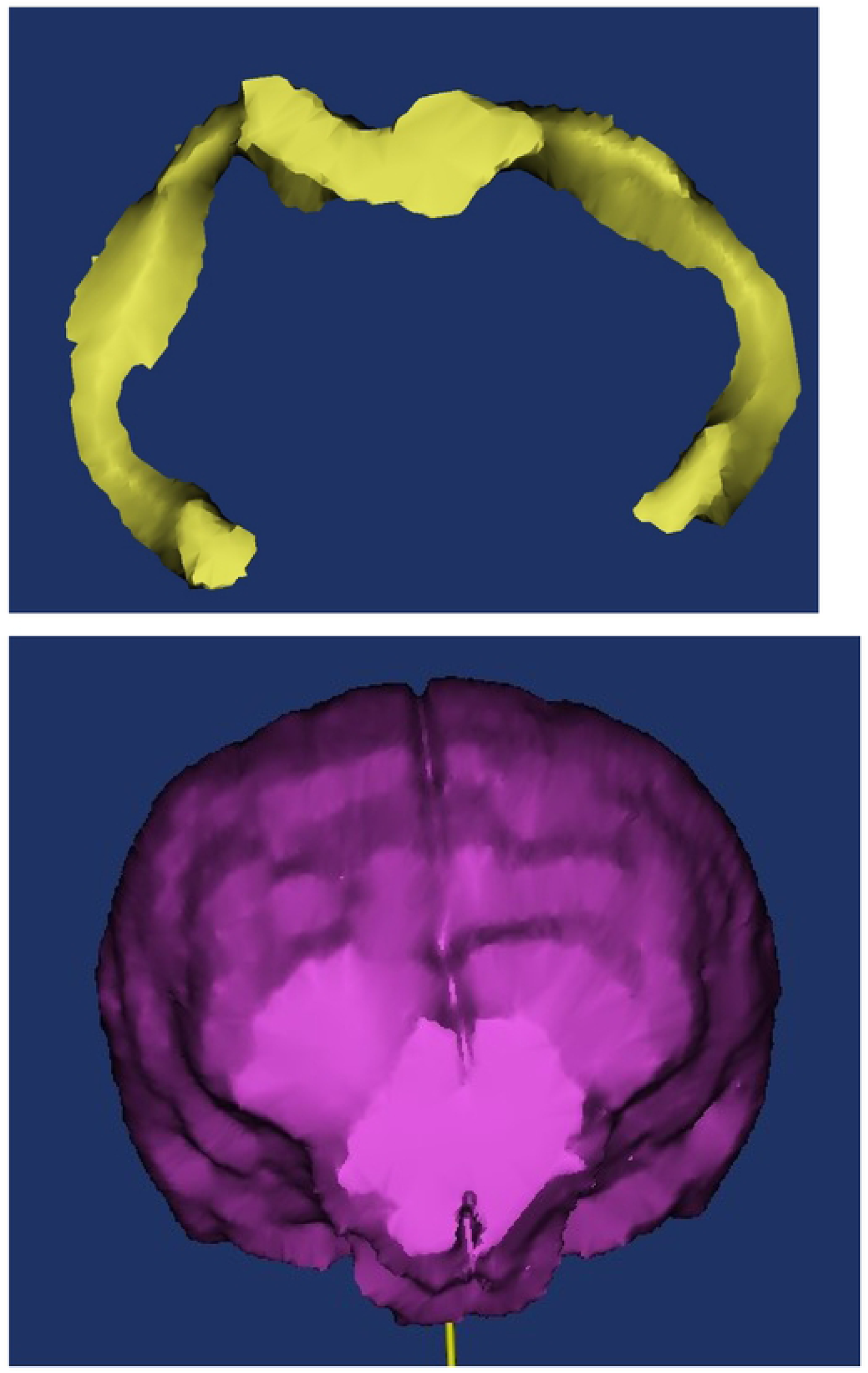
Examples of hippocampal volume (A) and total brain volume (B) procurement from MR images using Mimics^®^ software.

We then normalized total hippocampal volumes to total brain volume (rather than bodyweight) under the assumption that total brain volume would not change with CCD, and that total brain volume (but not bodyweight) remains unaffected by body condition (i.e. obesity, emaciation) according to the following equation: 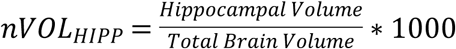

Because hippocampal volume represents a small percentage of total brain volume, we multiplied the volume ratio by 1000 to have more easily understood values.

### Statistical Analyses

We compared all continuous variables (i.e. normalized hippocampal volumes, age) between the CCD dogs and control dogs using Mann Whitney U Tests.

To assess for both intra and inter-observer variability in measurements, 20 patient scans were were re-measured by both observers (JS and MO), neither of whom were aware of the patient status (CCD vs control). Agreement between repeated measurements was examined using Limits of Agreement analysis.^15^

## Results

We included 16 dogs with CCD and 26 control dogs in the study. Dogs with CCD were older than control dogs (median age 13 yrs vs 11.5 yrs; P=0.0002); however, we could detect no negative correlation between advancing age and reduced hippocampal volume in either group (Figure 2). The CCD group comprised 2 Shih Tzus, 2 Springer spaniels and one each of Chihuahua, Miniature Poodle, Wheaten terrier, Labrador retriever, Tibetan terrier, Samoyed, Miniature Schnauzer, Cockapoo, German Shepherd, Shetland Sheepdog, Beagle and mixed breed. These consisted of 9 spayed females, 5 neutered and 2 intact males. The control group comprise 12 mixed breed dogs, 4 Chihuahuas and one each of Maltese, Boston terrier, Yorkshire terrier, Miniature Dachshund, Coonhound, Golden Retriever, West Highland White terrier, Beagle, Havanese and English Cocker Spaniel. These consisted of 13 spayed and 2 intact females, 7 neutered and 4 intact males.

**Figure 2.**
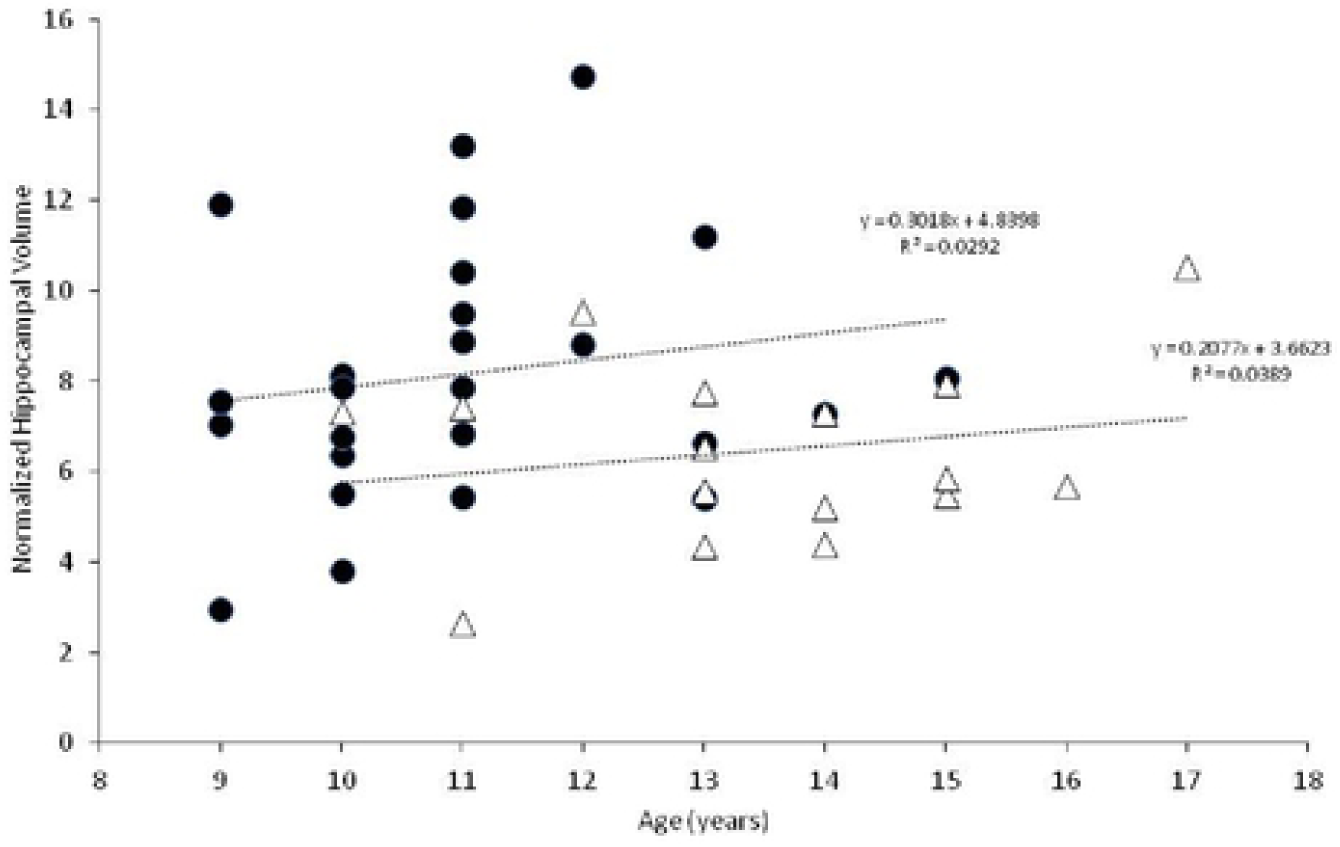
Normalized hippocampal volumes vs. age for controls (black circles) and CCD dogs (open triangles). Logistic regression analysis is provided for each group.

Dogs with CCD had smaller normalized hippocampal volumes than control dogs (median 6.2 vs 7.9; P=0.04; Figure 3).

**Figure 3.**
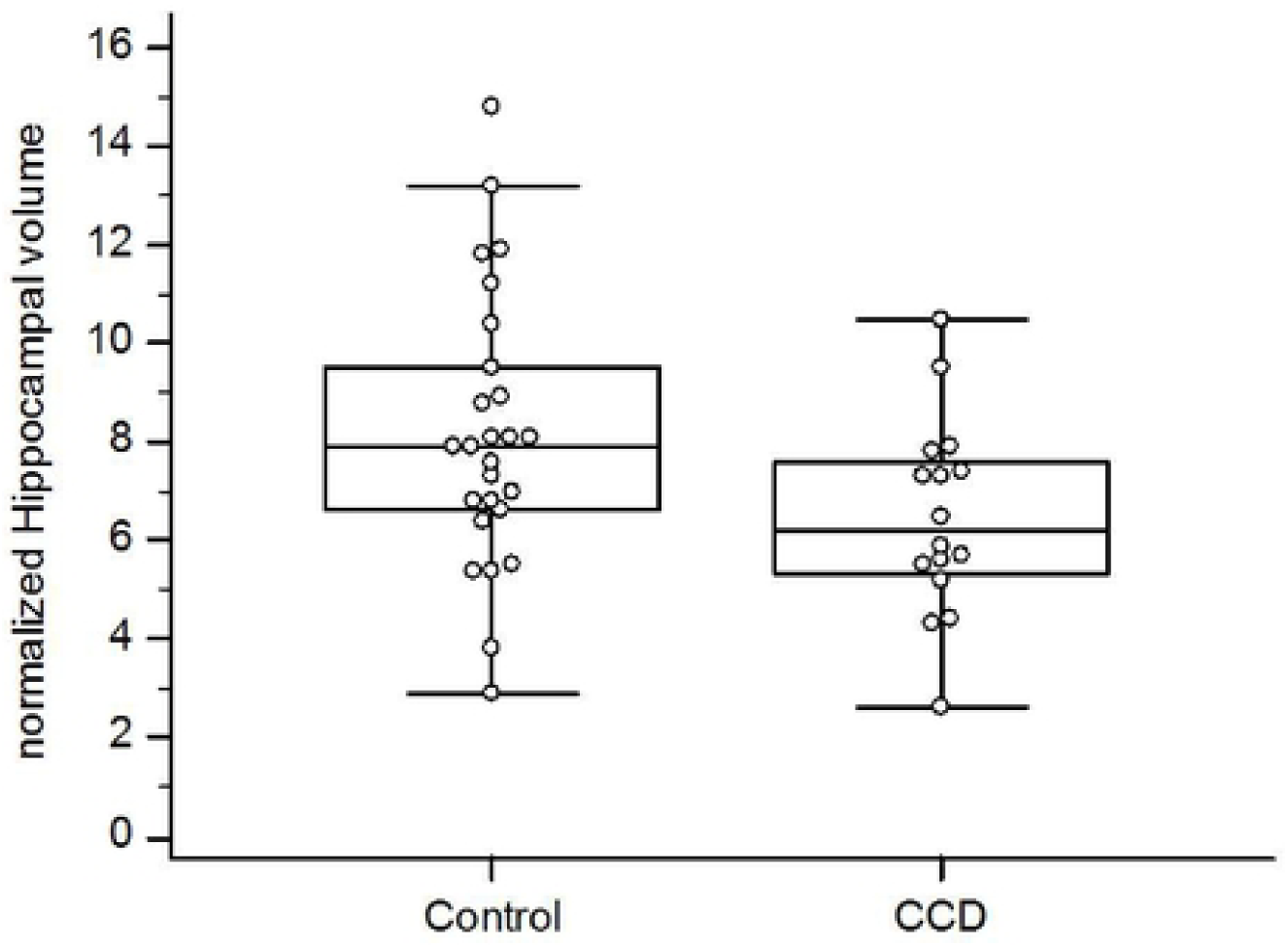
Box and whisker plots comparing normalized hippocampal volumes between aging control dogs and dogs with CCD.

## Discussion

Our study demonstrates that dogs afflicted with CCD have smaller hippocampal volumes compared to similarly aged control dogs without CCD. The differences were small, and showed a substantial degree of overlap between the two groups, suggesting that hippocampal volumetric measurements in dogs will be unable to discriminate between healthy and CCD dogs on an individual basis. However, our data suggest that hippocampal atrophy is not part of canine aging, as we could find no clear association between age and hippocampal volumes in either group.

Hippocampal atrophy is a key and early feature of human Alzheimer’s disease, and loss of hippocampal neurons and synapses is strongly associated with cognitive decline in that disorder.^3,4,16^ Hippocampal pathology has been documented in brains of dogs with CCD, but hippocampal atrophy has not been demonstrated for this disorder.^5^ In addition to hippocampal atrophy being a central pathophysiologic aspect of Alzheimer’s disease, the hippocampus is a source of neuronal stem and progenitor cells. Research into the roles of the hippocampus in Alzheimer’s disease pathogenesis and potential methods of positively affecting hippocampal function to treat Alzheimer’s disease patients is ongoing.^17-21^ CCD is considered a naturally-occurring canine analogue of human Alzheimer’s disease.^1,2,5,10,11^ Hippocampal-directed research in CCD patients may potentially benefit dogs with that disorder and human Alzheimer’s disease patients.

There are several limitations to this study, most of which are related to its retrospective nature. Although multiple institutions were involved in recruiting case material, the case numbers are still small. Also, the MR images evaluated were derived from multiple different machines, which could introduce some level of variability in the resultant data. We restricted case enrollment to dogs 9 yrs and older, in accordance with previous publications dealing with aging dogs.^10,23,24^ Although the median ages of our CCD and control groups was not large (11.5 yrs vs 13 yrs), it was statistically significant. A major hurdle in this investigation was locating control MRIs for comparison, most likely due to the low likelihood of dog owners pursuing brain MRIs for very old dogs without evidence of neurologic impairment. The possibility exists that the smaller hippocampal volumes in our CCD group were due to this group being older than the control dogs, vs a sequela to a degenerative brain disorder. The authors consider this unlikely for several reasons. Graphic representation (Figure 2) of hippocampal volumes vs age do not support a decreasing volume with aging for either the control or CCD groups. In addition, logistic regression analysis of these data also failed to discern a negative correlation between advancing age and hippocampal volume in either group. Age-related hippocampal atrophy has been documented to occur as an aging change in dogs, when young dogs are compared with older dogs.^23,24^ In one study, linear MRI measurements of the hippocampi normalized to brain height were compared between young (1-3 year old) and older (>10 year old) dogs with normal brain anatomy; a significant reduction of 2.64% was found between young and old dogs in that study.^23^ Although the results of our volumetric study of older dogs is not directly comparable to the results of the linear MRI study, the percentage difference between our two groups of dogs was 21.5%. In a study of laboratory Beagle dogs, hippocampal volumes were compared between young and old dogs using MRI. The older dogs were subdivided into two categories: old dogs (aged 8-11 years) and senior dogs (aged 12 years and older). Although hippocampal volume was shown to decrease when older dogs were compared to younger dogs (< 8yrs of age), there was no difference in hippocampal volume between the old and senior dog groups.^24^ In other words, age-related hippocampal atrophy appears not to progress dramatically as a non-specific aging change in normal dogs over 8 years of age, based on the results of the Beagle study.

Future MRI investigations into hippocampal atrophy in dogs with CCD would benefit from prospective investigations with larger case numbers, more closely age-matched controls, more consistent imaging procurement (i.e., one machine model), and more structurally detailed images (e.g., diffusion tensor imaging and tractography). Additionally, comparison of linear MRI measurements of hippocampal volumes between CCD patients and successfully aging dogs should be performed. Hopefully, investigations into hippocampal-related aspects of Alzheimer’s disease will benefit from CCD dogs as a disease model.

In conclusion, we demonstrated that dogs with CCD have significantly smaller total hippocampal volumes, as measured on MR images, compared with successfully aging controls. This finding may have implications in pathophysiologic and therapeutic research into hippocampal-associated aspects of CCD and Alzheimer’s disease.

